# Seroprevalence and risk factors for *Toxoplasma gondii* transmission in wild, domestic and companion animals in urban informal settlements from Salvador, Brazil

**DOI:** 10.1101/2025.07.01.662552

**Authors:** Leonela Bazan, Hernán Darío Argibay, Waléria Borges-Silva, Luís Fernando Pita Gondim, Thaís Auxiliadora dos Santos Mattos, Juliete Oliveira Santana, Eduardo Mendes da Silva, Michael Begon, Hussein Khalil, Federico Costa, Ianei de Oliveira Carneiro

## Abstract

*Toxoplasma gondii* is a globally neglected zoonotic parasite, particularly prevalent in socioeconomically vulnerable areas. Various animal species serve as reservoirs for *T. gondii* across different regions, including domestic cats, livestock, and a variety of wild and synanthropic animals. In urban areas, especially informal settlements, the close coexistence of humans, domestic animals, and wildlife may influence local transmission dynamics. This study evaluated the seroprevalence and associated risk factors for *T. gondii* infection in domestic and synanthropic animals from two low-income communities in Salvador, Brazil. A cross-sectional study was conducted in the neighborhoods of Marechal Rondon and Pau da Lima from October 2021 to February 2023. Blood samples were collected from domestic animals (288 dogs, 112 cats, 27 chickens, and six horses) and synanthropic species (54 brown rats and 75 big-eared opossums). Serological tests were performed using an indirect immunofluorescence antibody test. Questionnaires were used to collect environmental, demographic, and socioeconomic data from households where sampling took place. Generalized linear mixed models were applied to identify predictors of exposure. Seroprevalence was highest in chickens (66.7%), followed by dogs (37.2%), rats (24.1%), cats (22.3%), opossums (20%), and horses (16.7%). No significant factors were found to be associated with *T. gondii* seroprevalence in chickens, horses, rats, or opossums. Nevertheless, in dogs and cats, homemade diets increased the odds of infection by nearly six times compared to commercial feeding. Dogs from Pau da Lima were twice as likely to be infected as those from Marechal Rondon. These findings underscore the importance of promoting safe pet management, improving sanitation, and monitoring sentinel species to mitigate zoonotic risks in urban informal settlements.

**Author summary:** *Toxoplasma gondii* is a parasite that infects humans and animals worldwide, significantly impacting impoverished areas. In this study, we investigated the prevalence and factors associated with *T. gondii* infection in domestic and urban wildlife from two low-income communities in Salvador (BA, Brazil). We collected blood samples from dogs, cats, chickens, horses, rats, and opossums to test for antibodies against the parasite and analyzed environmental and lifestyle factors that might influence infection risk. Chickens showed the highest infection rates (67%), followed by dogs (37%), rats (24%), cats (22%), opossums (20%), and horses (17%). We found that homemade diets significantly increased the likelihood of infection in dogs and cats compared to commercial pet food.

Additionally, dogs from one neighborhood were twice as likely to be infected as those from the other, likely due to environmental conditions. These findings highlight how diet, environment, and urban living conditions affect the spread of *T. gondii*. By improving sanitation, promoting responsible pet care, and monitoring animals that share human environments, we can reduce the risk of this parasite in vulnerable communities.

## Introduction

*Toxoplasma gondii* (Nicolle and Manceaux, 1908) is an apicomplexan coccidian responsible for toxoplasmosis, a neglected zoonotic parasitic disease associated with poor living conditions (1,2). Approximately 30% of the global human population shows evidence of exposure to *T. gondii* (3); however, seroprevalence varies significantly worldwide (4). In highly endemic regions, such as Brazil, it can reach nearly 90% in specific demographic groups (5). *T. gondii* definitive hosts are members of the felid family, while all birds and mammals, including domestic and wild animals, serve as intermediate hosts (3,6). *T. gondii* undergoes sexual reproduction in the intestines of felids, with oocysts excreted in the faeces (7). Under suitable humidity and temperature, these oocysts become infective within 2-3 days and can persist in the environment, contaminating soil, water, and vegetation (8). Oocysts are highly resistant and, under suitable humidity and temperature conditions, can remain infective in soil for up to 18 months (9)and in water for 18 to 54 months under experimental conditions (8,10). A single felid can excrete millions of oocysts, leading to widespread contamination of soil, water, vegetables, gardens, and recreational areas (11). *T. gondii* transmission occurs mainly through ingesting contaminated food or water and congenital transmission from mother to fetus during pregnancy (12). Urban animals such as pets, poultry, rats, and opossums can serve as intermediate hosts for *T. gondii*, where the parasite multiplies asexually, forming tissue cysts in organs such as muscles and the brain. These intermediate hosts play a significant epidemiological role in human, animal, and environmental health. As they share the same environment and familiar sources of infection as humans, these animals can act as effective sentinels for monitoring *T. gondii* exposure (13–16).

As definitive hosts, domestic cats are the primary source of oocyst shedding in urban ecosystems (17). Cats are mostly infected during the first months of life, with prevalence increasing in those with street access or those fed homemade or raw meat (18). Other urban mammals and birds act as intermediate hosts and can become infected by ingesting water, soil, or food contaminated with oocysts and consuming infected animals (12). Due to their coprophagic habits, tendency to roll in cat feces, and close contact with their owners, pet dogs can indicate human contamination risk (19,20). Chickens are particularly effective indicators of oocyst contamination in soil, as they forage constantly and remain in close contact with the ground (21). Synanthropic rodents indicate environmental contamination by *T. gondii* oocysts and are a primary source of infection for definitive hosts, playing a significant role in ecological dissemination (22,23).

Given the technical challenges in directly quantifying oocysts in the environment, assessing the serological status of *T. gondii* in free-ranging urban animals is a valuable proxy for evaluating environmental contamination and associated epidemiological risks to human populations (16,24,25). Using urban animals as sentinels to monitor the spread of zoonotic pathogens is essential, as health risks are interconnected across species, and the emergence and persistence of these diseases are driven by complex, multidisciplinary factors (15). Examining demographic, social, and environmental factors associated with *T. gondii* exposure in sentinel animals enables the implementation of preventive measures to reduce pathogen exposure within communities (19). There is limited research on the prevalence and risk factors of *Toxoplasma gondii* infection in animal communities within informal urban settlements. Further research is needed to assess *T. gondii* epidemiology in local animal populations, in order to enhance understanding and improve infection control strategies in the region (26). In particular, little is known about the dynamics of *T. gondii* infection in chickens under natural conditions (27). This study presents a comprehensive analysis encompassing companion animals (cats and dogs), domestic species (chicken and horses), wildlife (opossums), and synanthropic species (rats). By including such a diverse range of species, our research provides valuable insights into the transmission dynamics and eco-epidemiology of *T. gondii* among animals living in these environments. This study aims to determine the seroprevalence and associated risk factors for *T. gondii* infection in domestic and synanthropic animals within informal urban settlements in Salvador (BA, Brazil), thereby contributing to the understanding of infection dynamics and supporting the development of targeted prevention strategies.

## Methodology

### Study area

This study was conducted in Salvador (Bahia, Brazil), located in the northeast of the country, with a population of 2,418,005 (IBGE, 2023). The eco-epidemiological research was conducted in two informal urban settlements in the Marechal Rondon and Pau da Lima neighborhoods, using polygons defined by previous epidemiological studies (28) (Figure 1). These communities were selected due to their shared characteristics typical of vulnerable urban areas: lack of urban planning, inadequate basic sanitation, and unfavorable socioeconomic conditions (29,30).

**Figure 1:**
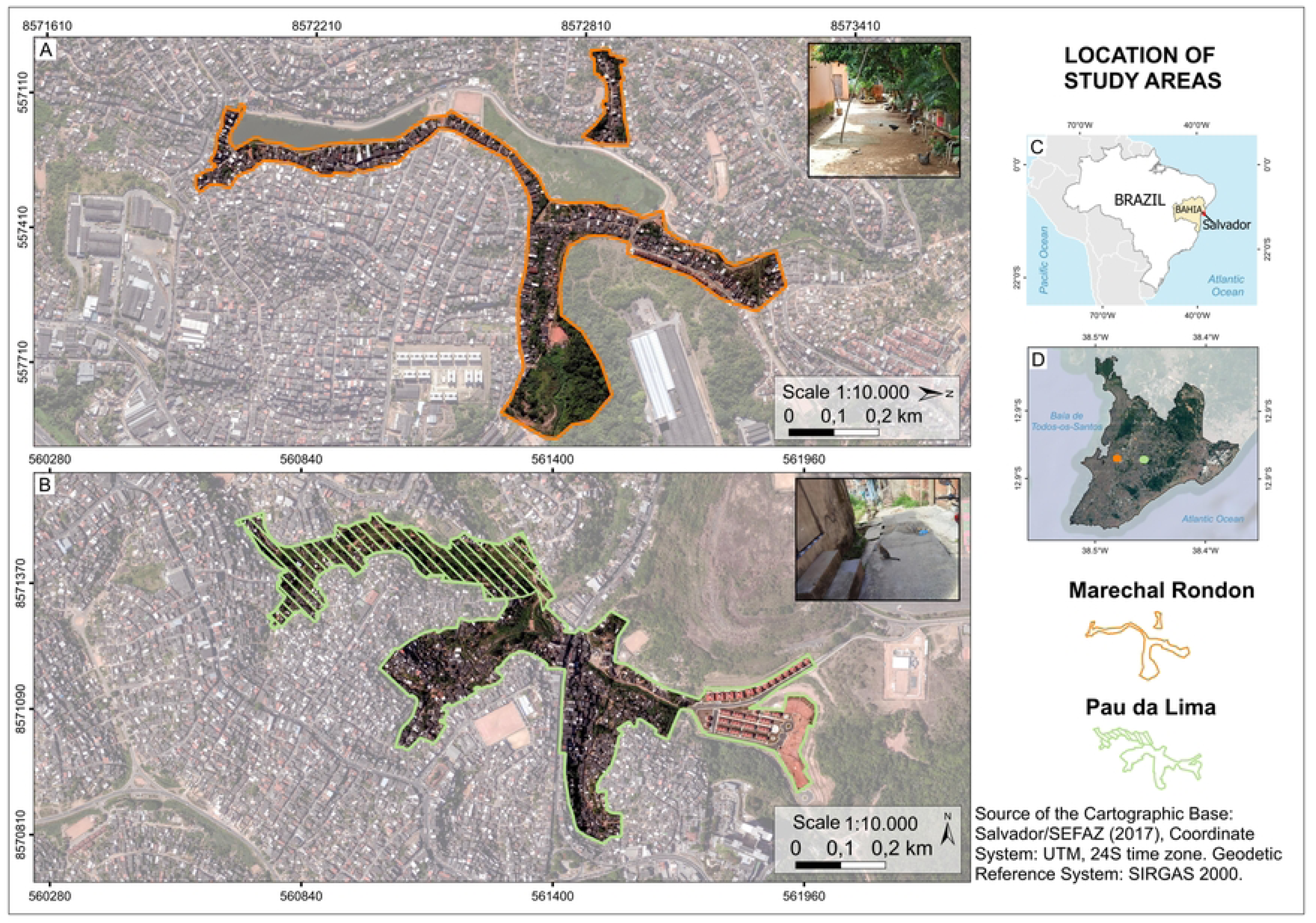
(A) The study area in the Marechal Rondon neighborhood is delineated by an orange line. (B) The study area in the Pau da Lima neighborhood is delineated by a green line. The area filled with diagonal lines represents the zone used for domestic animal sampling. (C) Map of Brazil showing the location of Salvador in the state of Bahia. (D) Map of Salvador (Bahia, Brazil) indicating the Marechal Rondon neighborhood with a green dot and Pau da Lima with an orange dot.

### Study design and sample collection

This research was approved by the Ethics Committee on the Use of Animals (CEUA) of the School of Veterinary Medicine and Animal Science at the Federal University of Bahia (protocol 07/2021), as well as by the Biodiversity Authorization and Information System (SISBIO, protocol 77314-1).

A cross-sectional study was conducted within a Community-Based Participatory Research (CBPR) project to collect samples from animals in communities between October 2021 and February 2023, parallel to a longitudinal epidemiological study in the same area (31). Samples were collected from domestic animals (dogs, cats, chickens, and horses), wild (opossum), and synanthropic animals (rats) for serological analysis.

For the domestic animals’ sample collection, all households within the polygons were identified with a unique code. Subsequently, all homes in the study area were visited, inviting domestic animal owners to participate in the research project. During the sampling period, the homes of residents who agreed to participate in the study and who signed a consent form were visited to allow the collection of biological samples from the animals. The animals were physically restrained to obtain general clinical data and to collect peripheral blood through cephalic or jugular venipuncture. Chemical restraint was selectively used for reactive cats (32). Following the collection of blood samples, a semi-structured questionnaire was applied to the owner to assess epidemiological variables related to the clinical history, demographic information about the animal, and socio-environmental conditions of the home.

To ensure an adequate representation of the community, 114 and 107 evenly spaced random points were selected in Marechal Rondon and Pau da Lima, respectively, for sampling wild and synanthropic animals (S1 Figure). Two Tomahawk live traps baited with sausage and pineapple were deployed at each point for four nights, resulting in 910 attempted trap-nights in Marechal Rondon and 856 in Pau da Lima. The traps were checked every morning, and their baits replaced when necessary. The traps were situated in peri-domestic settings, such as backyards, following the acquisition of permission from the residents, and in public areas. For each sampling point, a questionnaire was administered by field personnel to assess the environmental characteristics of the peri-domestic area. Following their capture, the rats and opossums were transported to a field laboratory, where they were taxonomically identified, their body measurements were taken, and blood samples were collected using sterile vacuum blood collection tubes. General clinical notes and overall physical condition were recorded. The captured opossums were anesthetized with ketamine (15mg/kg) and midazolam? (0.25mg/kg), administered intramuscularly (33) prior to biological sample collection. The rats received ketamine (25 mg/kg) with xylazine (3 mg/kg) via intramuscular injection, and after blood collection by cardiac puncture, thiopental (100 mg/kg) was administered intraperitoneally.

### Serological testing

Serum extraction was performed by centrifugation at 1811 x *g* for 15 minutes. Serum samples were aliquoted in 1.5 mL microtubes, identified, and stored at 20 °C until the serological test was performed. The detection of antibodies to *T. gondii* were detected using an indirect immunofluorescence antibody test (IFAT) with RH strain tachyzoites obtained through cell culture, with minor modifications (34). Sera were diluted in phosphate-buffered saline (PBS), and the cut-off point was 1:50. Positive and negative controls of each species were used for each reaction. For the detection of IgG anti-*T. gondii* antibodies, were used commercial anti-Cat IgG and anti-Rat IgG produced in goats (–Sigma-Aldrich®) anti-Dog IgG, anti-Horse IgG, and anti-Chicken IgY produced in rabbit (Sigma–Aldrich®) labeled with fluorescein isothiocyanate. The Zoonoses and Vector-borne Diseases Laboratory (DVZ, COVISA) produced the anti-opossum IgG in goats. The conjugates were diluted at 1:400 for dogs and chickens, 1:300 for cats, 1:100 for horses, and 1:50 for rats and opossums. These dilutions were determined based on preliminary tests using positive control samples.

### Statistical analysis

The seroprevalence was estimated as the percentage of individuals with a positive serological status on the IFAT, with a 95% confidence interval (CI). Prevalence differences among species were evaluated using the chi-square test. The positivity or negativity of each sample constituted the response variable used in the statistical analyses. All statistical analyses were performed using R Studio software (RStudio Team, 2020). To investigate the best model to explain the serological status of *T. gondii*, generalized linear mixed models (GLMM) were applied with lme4 package (35), with the household for domestic animals or collection point for wild/synanthropic species, as a random variable. Model selection was based on Akaike Information Criterion (AIC) values using the "MuMIn" package (36). The plausible models considered were those whose AIC difference was less than two compared to the best model. Among these, the final model was the most parsimonious, that is the one with the fewest explanatory variables. To analyze the collinearity between the variables, variance inflation factors were calculated, considering only those with a factor less than four. Spatial distribution of seropositive and seronegative animals was visually described by the construction of maps using the geographic coordinates of the collection points in the QGIS version 3.26.0 program (QGIS Development Team, 2009).

In this study, we included a range of explanatory variables to evaluate factors associated with *T. gondii* seroprevalence in domestic and synanthropic animals within urban communities. For domestic animals, variables related to individual characteristics and environmental conditions were assessed. These included neighborhood, sex, sterilization status, age, type of diet, shelter type, management practices, vaccination, deworming status, household crowding (residents per room), garbage disposal practices, garbage deposit, access to paved areas, wall material, ground slope, activity of the Zoonosis Control Center (CCZ), backyard paving, presence of the peri-domestic regions, and frequency of garbage collection. For dogs, we excluded collinear variables, resulting in a complete model that retained neighborhood, sex, sterilization status, age, type of diet, shelter type, management practices, vaccination, deworming, residents per room, garbage deposit, paved access, ground slope, CCZ activity, backyard paving, peri-domestic areas, and garbage collection. For cats, the final model also excluded collinear variables, leaving neighborhood, age, type of diet, shelter type, management practices, vaccination, residents per room, garbage disposal practices, garbage deposit, wall material, ground slope, backyard paving, and garbage collection. For synanthropic animals, we included neighborhood, sex, age, body condition, and vegetation coverage.

## Results

We collected blood samples from 288 dogs (*Canis lupus familiaris*), 112 cats (*Felis silvestris catus*), 27 chickens *(Gallus gallus domesticus*), six horses (*Equus ferus caballus*), 54 brown rats (*Rattus norvegicus*), and 75 big-eared opossums (*Didelphis aurita*). Of the total number of animals, 107/288 dogs (37.2%; 95% CI: 31.8 – 42.9), 25/112 cats (22.3%; 95% CI: 15.6 – 30.9), 18/27 chickens (66.7%; 95% CI: 47.8 – 81.4), 1/6 horses (16.7%; 95% CI: 0.42 – 64.1), 13/54 brown rats (24.1%; 95% CI: 14.6 – 37) and 15/75 big-eared opossums (20%; 95% CI: 12.5 – 30.4) were found to be seropositive for *T. gondii* (Table 1, Fig. 2, Fig. 3, S1 Table, S2 Table, S3 Table). A significant association was found between species and *T. gondii* seroprevalence status (Chi-square = 30.42, df = 5, *p* < 0.001), indicating that prevalence varied significantly among species.

**Figure 2:**
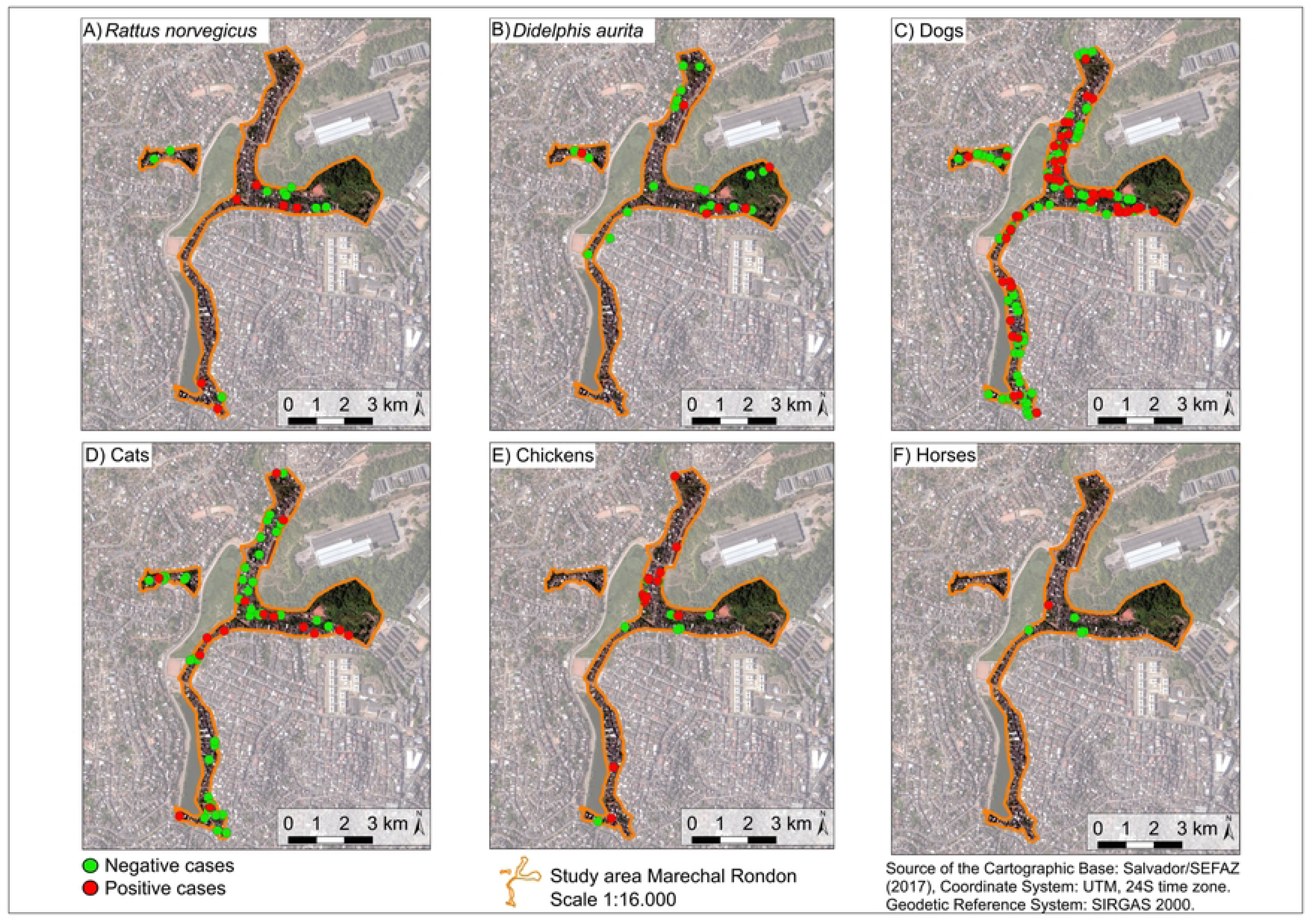
Spatial distribution of sampled individuals by species and frequency of *T. gondii*-seropositive animals in Marechal Rondon.

**Figure 3:**
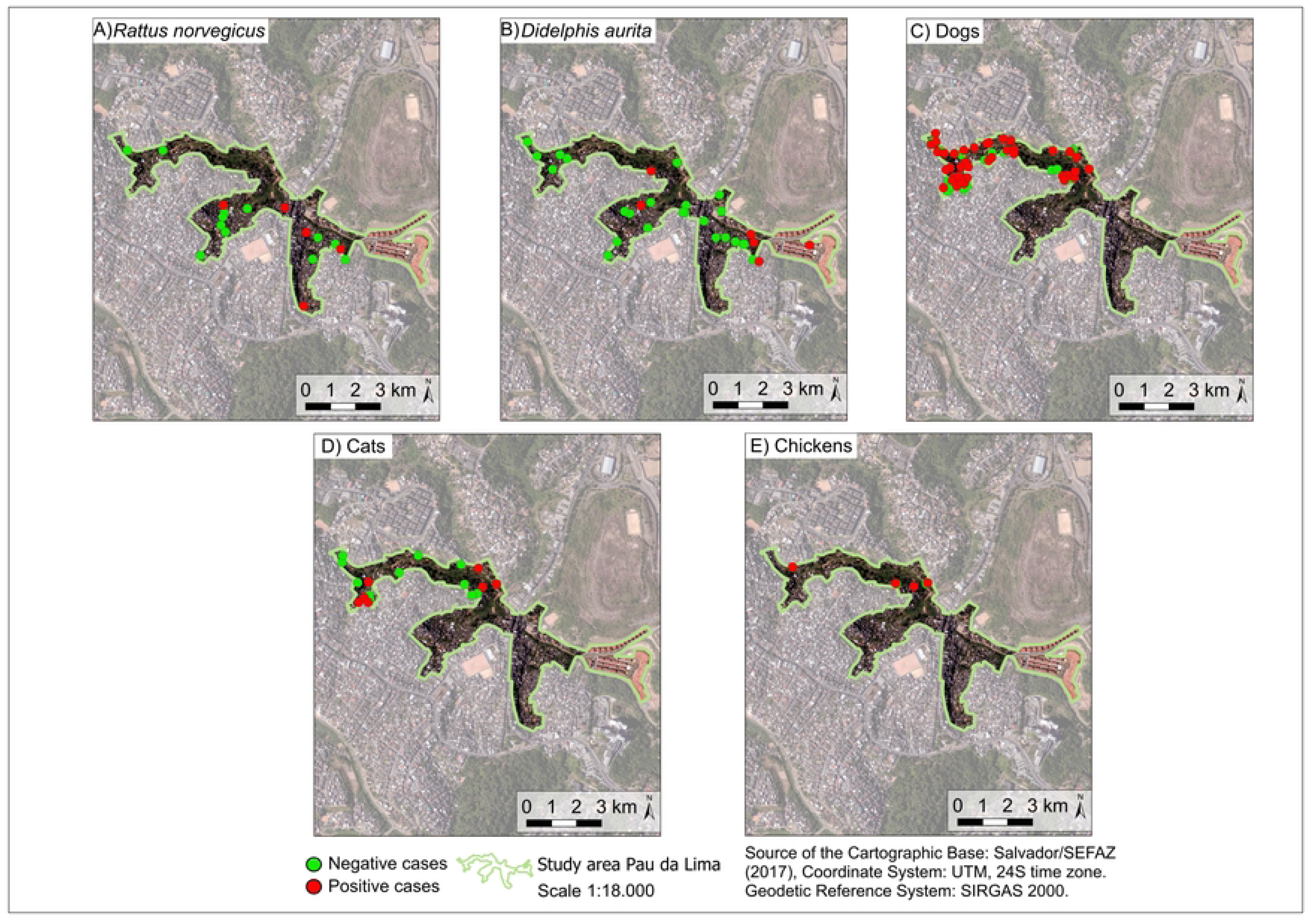
Spatial distribution of sampled individuals by species and frequency of *T. gondii*-seropositive animals in Pau da Lima.

**Table 1:** Frequency and seroprevalence of *Toxoplasma gondii* antibodies in animals from the Marechal Rondon and Pau da Lima neighborhoods.

| Specie and neighborhood | Total | Positives | Prevalence (%) | 95% CI |
| --- | --- | --- | --- | --- |
| Dogs |  |  |  |  |
| Marechal Rondon | 173 | 53 | 30.6 | 23.9 – 38.1 |
| Pau da Lima | 115 | 54 | 47.0 | 37.6 – 56.5 |
| Cats |  |  |  |  |
| Marechal Rondon | 86 | 18 | 20.9 | 12.9 – 31.0 |
| Pau da Lima | 26 | 7 | 26.9 | 11.6 – 47.8 |
| Chickens |  |  |  |  |
| Marechal Rondon | 21 | 14 | 66.7 | 43.0 – 85.4 |
| Pau da Lima | 6 | 4 | 66.7 | 22.3 – 95.7 |
| Horses |  |  |  |  |
| Marechal Rondon | 6 | 1 | 16.7 | 0.42 – 64.1 |
| Pau da Lima | 0 | 0 | - | - |
| Brown rats |  |  |  |  |
| Marechal Rondon | 25 | 7 | 28.0 | 12.1 – 49.4 |
| Pau da Lima | 29 | 6 | 20.7 | 8.0 – 39.7 |
| Big-eared opossums |  |  |  |  |
| Marechal Rondon | 36 | 8 | 22.2 | 10.1 – 39.1 |
| Pau da Lima | 39 | 7 | 17.9 | 7.5 – 33.5 |

In the final multiple regression model, the type of diet and neighborhood were associated with *T. gondii* seropositivity in dogs. In contrast, only the type of diet was associated with *T. gondii* seropositivity in cats. Dogs fed homemade food were almost six times as likely to be exposed to *T. gondii* (OR: 5.60; 95% CI 2.72 - 11.95) compared to dogs fed commercial food. Dogs fed a mixed diet were also found to be almost three times as likely to be exposed to *T. gondii* (OR: 2.82; 95% CI 1.50 - 5.52) compared to dogs fed commercial food. Dogs in Pau da Lima exhibited a twofold increased likelihood of exposure to *T. gondii* (OR: 1.95, 95% CI 1.17 - 3.26) compared to dogs in Marechal Rondon. In cats, consumption of homemade foods increased the probability of infection by *T. gondii* by fivefold (OR: 5; 95% IC 0.84 – 30.03), while consumption of mixed foods increased the likelihood of infection by *T. gondii* by twice (OR: 2.08; 95% IC 0.78 - 5.48) (Table 2).

**Table 2:**
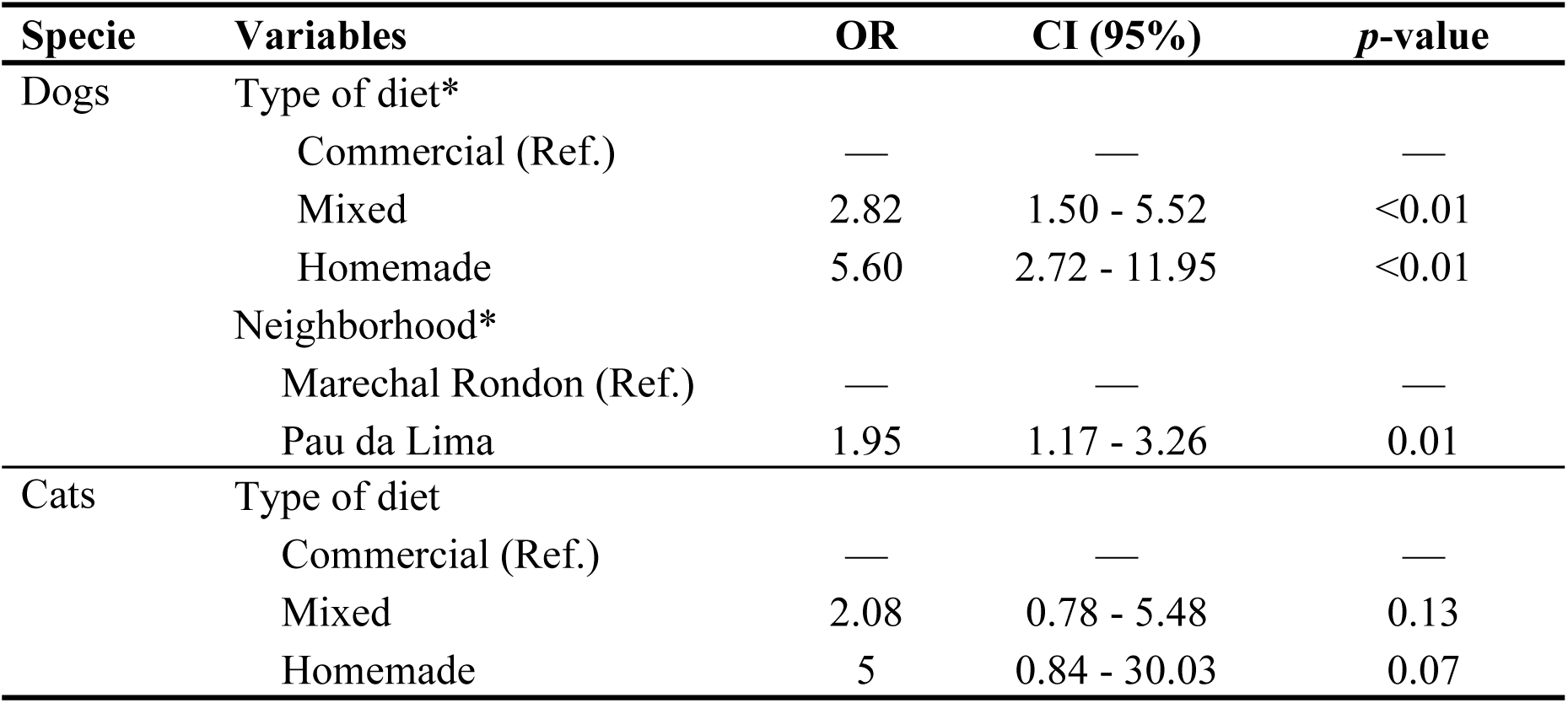
Multivariate analysis of the associated factors of *Toxoplasma gondii* seroprevalence in dogs and cats from Marechal Rondon and Pau da Lima, Salvador, BA, Brazil. OR: odds ratios; CI: 95% confidence interval; * Categories with a significant effect (p-value < 0.05).

## Discussion

In this study, the highest seroprevalence of *T. gondii* was observed in chickens (66.7%), followed by dogs (37.2%), brown rats (24.1%), cats (22.3%), big-eared opossums (20%), and horses (16.7%). Diet and neighborhood were identified as significant factors influencing *T. gondii* exposure in dogs, whereas only diet type was linked to seropositivity in cats. Our findings revealed that animals fed homemade food and dogs from Pau da Lima were at greater risk of infection. These results highlight the critical role of dietary habits in assessing *T. gondii* exposure in domestic animals and emphasize their potential implications for public health.

The type of diet was identified as a common factor associated with *T. gondii* exposure in dogs and cats. Our findings show that dogs and cats fed homemade food are more likely to be exposed to *T. gondii* than those fed commercial pet food. Consequently, feeding pets with commercial pet food is a protective factor against exposure to this pathogen. This observation aligns with the findings in São Paulo state that homemade feeding increases the odds of *T. gondii* exposure in dogs (37,38) and cats (39). Ingesting food contaminated with oocysts or tissue cysts is the primary source of *T. gondii* infection in animals and humans (6). As such, providing homemade food to pets may serve as a potential source of infection and contribute to the spread of *T. gondii* (37,40). Additionally, pets fed exclusively on commercial food are generally better nourished and less susceptible to infections (41,42).

It is also noteworthy that feeding pets only with commercial pet food may reflect better management practices by their guardians, such as not allowing dogs to roam freely, maintaining regular vaccination and deworming schedules, among others (43). These factors further reduce the likelihood of *T. gondii* exposure in the environment. Furthermore, dogs from Pau da Lima were twice as likely to be exposed to *T. gondii* than those from Marechal Rondon. The higher seroprevalence observed in Pau da Lima may be associated with socio-environmental factors that increase dogs’ exposure to the pathogen. Variations could also influence differences between neighborhoods in the knowledge, attitudes, and practices of pet guardians (44), which warrants further investigation in future research.

The seroprevalence of *T. gondii* in dogs in both neighborhoods was lower than the global seroprevalence of Brazil mentioned in Dubey *et al*. (2020), which is 70% (18). This discrepancy is likely because many studies cited used a lower cut-off point than the one employed in our research, potentially leading to inflated seroprevalence rates. The seroprevalence in Marechal Rondon matched the 34% reported in dogs in Rio de Janeiro that presented for veterinary care, encompassing general check-ups and toxoplasmosis diagnosis (45). Furthermore, it was comparable to the 30.7% prevalence seen in dogs from Curitiba that frequented densely occupied public spaces such as bus stations and parks (46). The seroprevalence in Pau da Lima was similar to the 48% reported in domiciled dogs on Fernando de Noronha Island (47) and dogs from urban informal settlements in Jataizinho, Paraná (48). However, it was lower than the 70.5% reported in partly-domiciled dogs from Londrina, Paraná (49), who employed a lower cut-off point (1:16) than ours, which may have led to a higher number of positive samples, and higher than the 9.5% reported in domiciled dogs from Garanhuns, Pernambuco (50).

The seroprevalence of *T. gondii* in cats in our study was comparable to the 21% observed in domiciled cats from Belém, Pará (51) and the 25% reported in cats from rural villages in the semi-arid region of Northeastern Brazil (52). In contrast, it was lower than the global seroprevalence of 35% (53) and the 50% reported in pet cats of pregnant women attending healthcare services in Ilhéus, Bahia (54).

The *T. gondii* seroprevalence in the chickens tested was higher than the global prevalence of 30% (55), similar to the 71% reported in free-range chickens from Minas Gerais (56). Backyard chickens are considered a potential source for spreading this pathogen, as they are often raised for egg and meat consumption. Chickens play a significant role in the epidemiology of *T. gondii*, arguably more so than rodents, due to their clinical resistance to the parasite and the ability of cats fed naturally infected chicken tissues to shed millions of oocysts (57). Poultry are an ideal sentinel species for monitoring environmental contamination with *T. gondii*, as their ground-feeding behavior exposes them directly to oocysts (21,27). In urban informal settlements, chickens are frequently slaughtered at home or in unsupervised facilities, with viscera often left for scavengers or improperly discarded. *T. gondii* infection may occur if hygiene measures, after handling or cooking poultry, are not strictly followed. However, comprehensive risk assessment studies addressing this issue remain limited.

The worldwide prevalence of *T. gondii* antibodies in brown rats is 6% and in South America is 18% (22), lower than the overall prevalence observed in this study (24%). Due to their feeding behaviour that predominantly facilitates oral transmission of sporulated oocysts within the environment, synanthropic rodents can be regarded as indicative of environmental oocyst contamination. Consequently, the finding of *T. gondii* in rat populations might reflect the dissemination of the parasitic environmental phase within a specific geographical area (23,58). Rats are recognized as reservoirs and carriers of the disease, serving as the primary source of cat infection (59). The *T. gondii* seroprevalence in big-eared opossums in this study was similar to the 22.7% reported in *Didelphis aurita* and *Didelphis albiventris* from urban areas in São Paulo state (60), and higher than the 5.5% reported in *Didelphis albiventris* also from São Paulo state (61). Big-eared opossums can become infected by the consumption of infected rats (62,63); in addition, they can contribute to controlling rat populations in communities. Hunting opossums in peridomestic environments for human consumption or as food for domestic animals remains a significant factor that substantially increases exposure to zoonotic pathogens, including *T. gondii* (64,65).

Due to its cross-sectional design and the use of serological techniques to detect antibodies, it was impossible to determine when the animals were exposed to *T. gondii*. Additionally, the sample size was not balanced across species or between neighborhoods, which may limit the accuracy for comparing the seroprevalence and finding an explanatory model for all species. This variation in sampling could influence the generalizability of findings and the robustness of statistical associations drawn from the data.

After the COVID-19 pandemic, studying zoonotic diseases in historically neglected communities has become an urgent priority, particularly in the context of significant social and health inequalities in Brazil and around the world. Therefore, this study provides insights into the eco-epidemiology of *T. gondii* in urban animals, which could serve as sentinels for environmental contamination in vulnerable neighborhoods. The seroprevalence of *T. gondii* antibodies indicates exposure to the parasite and highlights a potential risk of infection within these communities. The finding that diet was identified as the main factor associated with *T. gondii* exposure in both dogs and cats reinforces the importance of promoting educational initiatives and campaigns in these neighborhoods to inform residents about the risks of infection for both animals and humans. The findings of this study are crucial for understanding the eco-epidemiological dynamics of *T. gondii* in vulnerable communities and for shaping future public policies on environmental sanitation, as well as communication and environmental education programs in these areas.

## Availability of data and materials

The datasets generated and analyzed during the current study are available at https://zenodo.org/records/15091116.

## Acknowledgements

We extend our heartfelt gratitude to the Marechal Rondon and Pau da Lima community residents for their trust and cooperation in allowing us to collect samples from their animals. We would also like to thank all Ecology and Environment team members of the Building Healthy Communities in Brazilian Urban Slums (CASA) project for their dedication and contributions to this study. Furthermore, we are grateful to the Zoonoses and Vector-borne Diseases Laboratory (DVZ, COVISA) in São Paulo (SP, Brazil). Finally, we thank the Coordination for the Improvement of Higher Education Personnel (CAPES) of the Ministry of Education of Brazil for a MSc scholarship.

